# Noradrenergic administration improves cognitive flexibility after permanent damage to glutamatergic neurons in rat mediodorsal thalamus or thalamic nucleus reuniens

**DOI:** 10.64898/2026.02.16.706106

**Authors:** Jennifer J. Hamilton, Lisa Berriman, Shae Harrison-Best, John C. Dalrymple-Alford, Anna S. Mitchell

## Abstract

Cognitive flexibility, switching behaviour responses to changing task demands, is classically attributed to the prefrontal cortex. Prefrontal thalamocortical circuits from mediodorsal thalamus (MD) or thalamic nucleus reuniens (RE) are altered in neurological conditions with cognitive flexibility deficits. Interventions targeting thalamocortical interactions may offer therapeutic benefits. Using the attentional set-shifting task, we showed in rats that permanent lesions of MD or RE glutamatergic neurons caused dissociable cognitive flexibility deficits and that the selective ⍺2-adrenoceptor antagonist, atipamezole improved cognitive flexibility. RE damaged rats were impaired on the first of three intradimensional shift (ID) subtasks involving responding to novel stimuli pairings linked to the sensory dimension that reliably predicts reward. In contrast, MD damaged rats had intact ID performance but were impaired in the extradimensional shift (ED) subtask, whereby after task demands change rats are required to make a rapid shift in attending to and adapting choice responses to novel stimuli pairings linked to the previously irrelevant sensory dimension. RE damage did not disrupt ED performance. Intraperitoneal injections of atipamezole (1mg/kg), given 30-min prior to retesting with novel stimuli pairings improved cognitive flexibility despite permanent thalamic damage. VGluT2 labelling in the anterior cingulate cortex indicated neural network downstream effects with MD damage producing increased labelling relative to Sham operated controls, while RE damage showed decreased labelling. These findings demonstrate the dissociable influence of MD and RE during cognitive flexibility and suggest the use of noradrenergic regulation as a potential therapeutic option to reduce this cognitive dysfunction.

## Introduction

Cognitive flexibility, the ability to keep track of current task demands and switch choice responding rapidly in response to change, is dysfunctional in many neurological conditions including stroke, dementia and Parkinson’s disease, as well as neuropsychiatric conditions such as anxiety, depression, obsessive compulsive disorder and schizophrenia [1–5]. Both the mediodorsal thalamus (MD) and thalamic nucleus reuniens (RE) support cognitive flexibility via processes linked to attention, learning, memory, and decision-making through their reciprocal connections with the prefrontal cortex (PFC) and different regions of the medial temporal lobe [1–3,6–17].

Analogous attentional set-shifting tasks have been established in humans [18–20], monkeys [21–24] and rats [25]. In the rat version, the task first tests the ability to learn and maintain responding to stimuli linked to one sensory dimension (olfaction or touch) across new stimulus pairings (e.g., odours), requiring an intradimensional (ID) shift. The rat is then required to make an extradimensional (ED) shift to the previously irrelevant sensory dimension (e.g., now digging media) for reward. Rats, monkeys and humans with PFC damage have deficits in the ED shift, while rats with anterior thalamic damage or anterior cingulate cortex (ACC) chemogenetic disruption have deficits in ID shifts [25–28]. Thalamocortical interactions are also crucial for cognitive flexibility in the rat as permanent glutamatergic neuronal damage in the MD in rats also causes ED shift deficits [9], whereas RE lesions cause deficits when responding to the ID shift [13].

Noradrenergic (NA) efferent pathways from the brainstem innervate the thalamus including MD, RE, in addition to pathways coursing to forebrain structures, namely the mPFC and hippocampus from the locus coeruleus [29]. NA innervation is altered in various neurological conditions, including Parkinson’s Disease, Alzheimer’s Disease and schizophrenia [30,31]. Centrally, ⍺2-adrenoceptors are located throughout the brain and mediate the modulatory actions of norepinephrine [32,33] and blocking these ⍺2-adrenoceptors using atipamezole promotes the release of norepinephrine [34–36] and improves cognition including cognitive flexibility in humans and animal models [30,31].

In rats, improved cognitive flexibility was demonstrated after permanent damage to the mPFC and treatment with intraperitoneal (i.p.) injections of atipamezole [26], while manipulations to the locus coeruleus or deafferentation of noradrenergic pathways in the mPFC impair cognitive flexibility [37–39]. However, we do not know if deficits in cognitive flexibility after permanent damage in either MD or RE thalamic nuclei can be restored by administration of atipamezole. The PFC, MD, and RE are all affected in the early- and mid-phases of most neuropsychiatric and neurodegenerative conditions, so understanding the benefits of manipulations to the noradrenergic system after thalamic changes may provide therapeutic insights for NA treatments to help recover cognitive flexibility.

Here, we tested rats with permanent damage to MD or RE caused by a glutamatergic excitotoxin or Sham controls to identify cognitive flexibility deficits in the attentional set-shifting task (IDED). Given RE damage previously caused deficits in acquiring a single ID shift [13], we assessed using three consecutive ID shifts to determine whether RE also supports implementation across two additional ID shifts [15,40]. We found dissociable deficits with RE damage affecting the first ID shift only, and MD damage affecting the ED shift only. Next, the rats were retested on the IDED task with novel stimuli pairings for the two sensory dimensions and received i.p. injections of atipamezole 30-min prior to testing. We found atipamezole improved cognitive flexibility in all three groups of rats including those with permanent MD or RE damage.

## Materials and Methods

Detailed materials and methods available in Supplementary Materials.

### Animals

Twenty-four male Piebald Virol Glaxo cArc (315-415g) rats were housed four per cage with lights on a reverse 12-hour light-dark cycle. Rats were maintained at 85% of free-feeding body weight (12-18g/rat/day food) with *ad libitum* access to water. The University of Canterbury Animal Ethics Committee approved all experimental procedures (2022/14R).

### Neurosurgery to create permanent MD or RE thalamic damage

Rats were assigned to one of three lesion groups: MD, N=8; RE, N=8; Sham, N=8. Permanent bilateral MD or RE excitotoxic (NMDA) lesions were made while rats were deeply anaesthetised and secured in a Kopf stereotaxic frame using the following coordinates, MD: AP=-3.75mm anterior and -3.95mm posterior (opposite hemisphere), ML=±0.7mm, DV=-5.70mm, 20µl per infusion site; RE: 10° angle, AP=-2.3mm, ML=±1.1mm, DV=-7.0mm, 0.12µl per infusion site.

### IDED attentional set-shifting task

For this experiment, the attentional set shifting IDED task was run twice (IDED1 and IDED2; apparatus in Fig. 1A) using 8 different subtasks (Fig. 1B): simple discrimination (SD); concurrent discrimination and its reversal (CD, CDrev); intradimensional shift (ID1-3); extradimensional shift and its reversal (ED, EDrev). Prior to running in each test session, each rat received an i.p. injection (IDED1=saline 1mg/kg; IDED2=atipamezole 1mg/kg, Abcam; [26]) and was placed into a holding cage for 30-min. IDED2 used the same stimuli as IDED1 but the pairings and rewarded stimulus were shuffled so all pairings were novel. Open field exploration was measured before and after IDED2 to compare activity levels after atipamezole injection.

**Figure 1.**
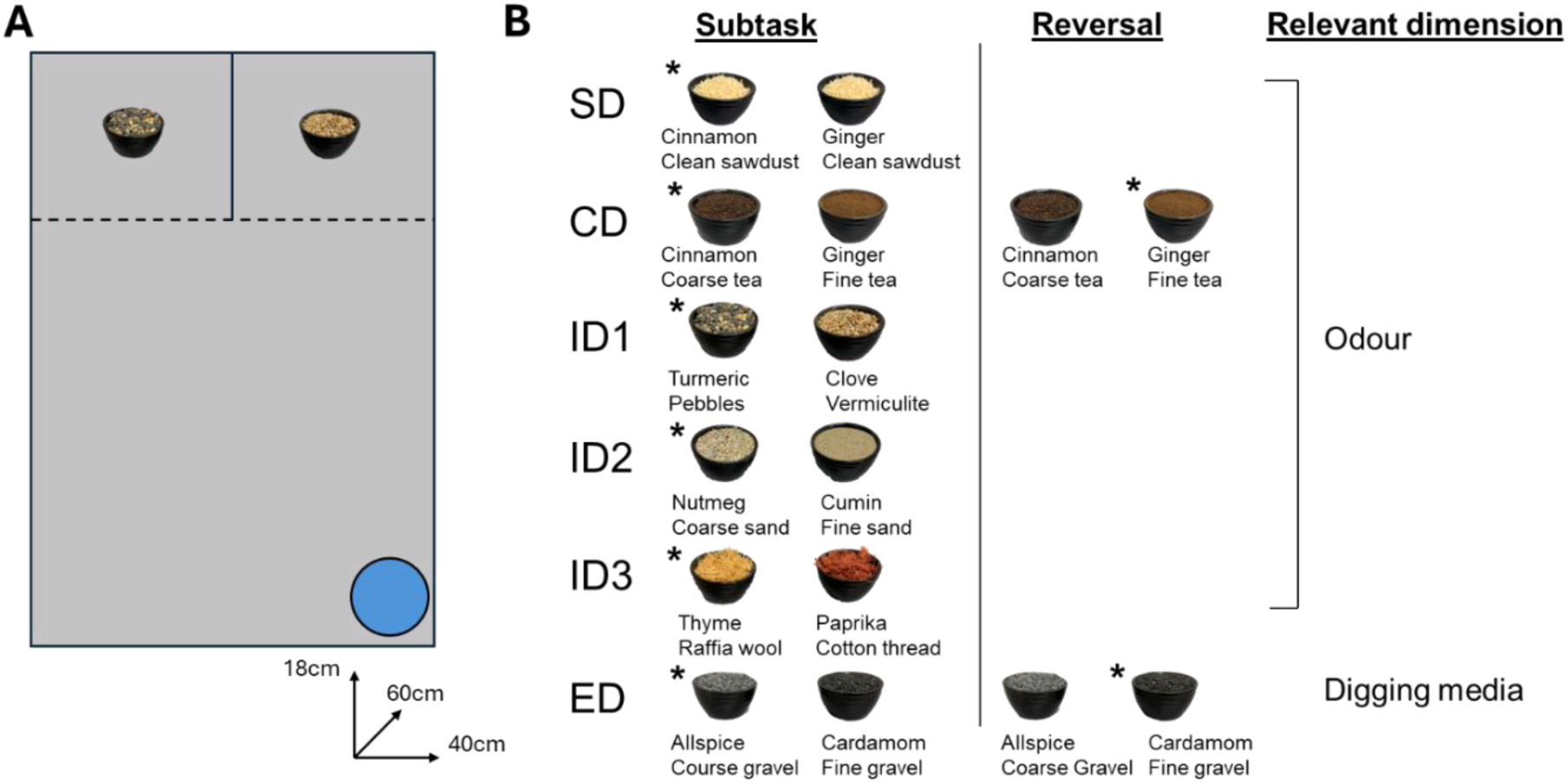
**A.** Schematic of the ID/ED attentional set-shifting apparatus. Access to two identical choice compartments was controlled by clear Perspex doors (dotted line). Blue circle = small glass jar to provide ad libitum water throughout testing. The black ceramic digging bowls were filled with a combination of odours and digging media. **B.** Stimulus pairing examples for each subtask of IDED test sessions. * = Correct odour/digging media (relevant dimension, stimuli and left/right were counterbalanced across individual rats), SD = simple discrimination, CD = concurrent discrimination, ID = intradimensional shift, ED = extradimensional shift.

### Histology

#### Perfusion and tissue collection

After completion of the experiment, deeply anaesthetised rats were perfused transcardially with saline and 4% paraformaldehyde (PFA) in 0.1M phosphate buffer. Coronal 40µm brain sections were collected using a freezing stage microtome (Bright Instruments, UK).

#### NeuN lesion verification

Per Hamilton & Dalrymple-Alford [41], sections throughout the thalamus were stained for NeuN (antibodies: anti-NeuN, 1:500, MAB377; Millipore, California USA; biotinylated goat anti-mouse, 1:1000 BP-9200-50: Vector Laboratories, California USA; diaminobenzidine, DAB 0.05%; Sigma). Sections were photographed at 5x objective on a Leica DM6B upright microscope and DFC7000T camera (Leica Microsystems, Germany). Image J analysis software (NIH, USA) was used to determine the percentage of lesion damage. Acceptable lesions were defined as at least 60% damage in the target thalamic structure.

#### VGluT2 immunofluorescence

mPFC sections were stained for VGluT2 (antibodies: anti-vGlut2, 1:1000; AB216463, Abcam, NZ; AlexaFluor488 goat anti-rabbit, 1:500, Invitrogen, Oregon, USA; DAPI Vectashield, Vector Laboratories, Burlingame, CA, USA). Sections were photographed at 40x objective with excitation from the L5 (green; Leica) and UV ‘A’ (blue; Leica) filters. Automated counts of the VGluT2 puncta were obtained through ImageJ.

#### Statistical analysis

Individual repeated measures ANOVAs evaluated the number of trials to criterion for each rat for each of the 8 subtasks with Lesion Group as the between subject factor for the IDED1 and IDED2. In IDED1, two rats did not complete the EDrev subtask due to time restrictions, therefore only seven subtasks were examined.

Due to the complex nature of the interaction between Subtask and Group, subsequent analyses were run. Based on *a priori* hypotheses of expected deficits following MD or RE damage, these were: Cluster 1=SD, CD, CDrev; Cluster 2=ID1, ID2, ID3; Cluster 3=ID3, ED; and ED subtask. Subtask clusters 1, 2 and 3 were analysed using repeated measures ANOVAs (e.g., [SD, CD, CDrev] x Group) with post-hoc analyses for relevant Cluster x Group effects (2-way ANOVA). ED subtask was analysed using a univariate ANOVA with pairwise comparisons. Across Drug Session [IDED1 vs IDED2] x Group, repeated measures ANOVAs compared trials to criterion for Cluster 1, 2 and the ED subtask and the Shift Cost, which used a difference score (ED minus ID3) to compare the trials to criterion of ID3 with ED. Finally, a repeated measures ANOVA compared differences between Groups of second entry choice responses made for all subtasks combined as a measure of rapid adaptability in choice responding.

## Results

### Assessment of MD and RE damage

NeuN quantification led to the exclusion of two rats, one MD and one RE (>40% target neuronal sparing). One Sham rat was excluded due to health concerns prior to completion of the experiment. Final group numbers were Sham controls, n=7, MD damage, n=7, RE damage, n=7. For MD rats included in the final analyses, the median damage to medial and central MD subdivisions was 79% (range 60-84%) while damage in the lateral MD subdivision averaged at 53% (see Fig. 2A). For RE rats, the median damage was 79% (range 70-93%; see Fig. 2A). In all cases, the other thalamic target was spared with minimal neuronal loss (no more than 10-20%). The central median thalamic region, between the MD and RE, also showed minimal cell loss.

**Figure 2.**
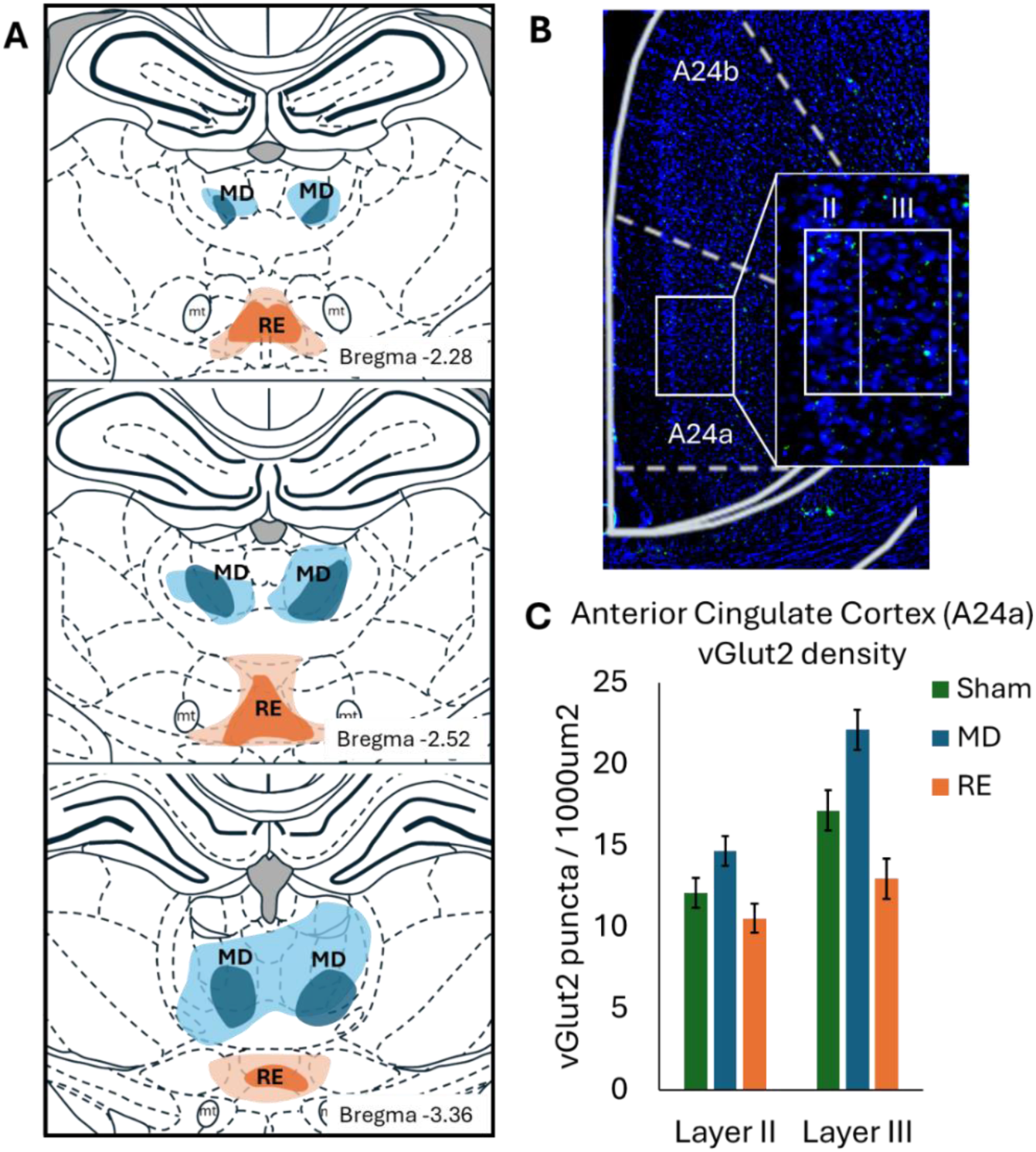
**A.** Schematic representation of the included smallest (dark blue = MD, dark orange = RE) and largest (light blue = MD, light orange = RE) NMDA neurotoxin damage at three different anteroposterior locations of the thalamus relative to bregma (mm). MD = mediodorsal thalamus, mt = mammillothalamic tract, RE = thalamic nucleus reuniens. **B.** Photomicrograph (40x magnification) images of DAPI (blue) and VGluT2 (green) in Layer II and Layer III of the anterior cingulate cortex region, A24a, approximately 1.28mm from Bregma. **C.** Mean (+/-SEM) for VGluT2 puncta density (per 1000um2) in the anterior cingulate area A24a Layers II and III for the three experimental groups: Sham control neurosurgery n=7; excitotoxic NMDA infusions into the mediodorsal (MD) thalamic nuclei, n=7 or into the thalamic nucleus reuniens (RE) n=7.

### VGluT2 expression in anterior cingulate (A24a)

As expected, there was no interaction effect for VGluT2 puncta density (Group x Layer *F*(2,18)=2.79, *p*=.08), but significant main effects for Group and for Layer were revealed (Fig. 2B&C). VGluT2 density was highest in MD rats, intermediate in Sham rats and lowest in RE rats (Group main effect, *F*(2,18)=18.02, *p*<.001; pairwise comparisons: Sham vs MD, *p*=.003; Sham vs RE, *p*=.01; MD vs RE, *p*<.001). Higher density of VGluT2 was present in Layer III compared to Layer II (*F*(1,18)=33.22, *p*<.001).

### Group effects on overall IDED1

Analysis of the first 7 subtasks (SD-ED) on IDED1 revealed a significant Subtask x Group interaction ([order 4], *F*(2,18)=9.74, *p*=.00). Rats with either MD (Sham vs MD, *p*=.006) or RE damage (Sham vs RE, *p*=.003; Fig. 3A) required more trials to criterion than Sham control rats. Trials to criterion for MD or RE rats did not differ across these subtasks (*p*=.69). There was also a Group main effect, *F*(2,18)=7.28, *p*=.005, and a significant Subtask main effect ([quad] *F*(1,18)=4.56, *p*=.04). These effects are further described below.

**Figure 3.**
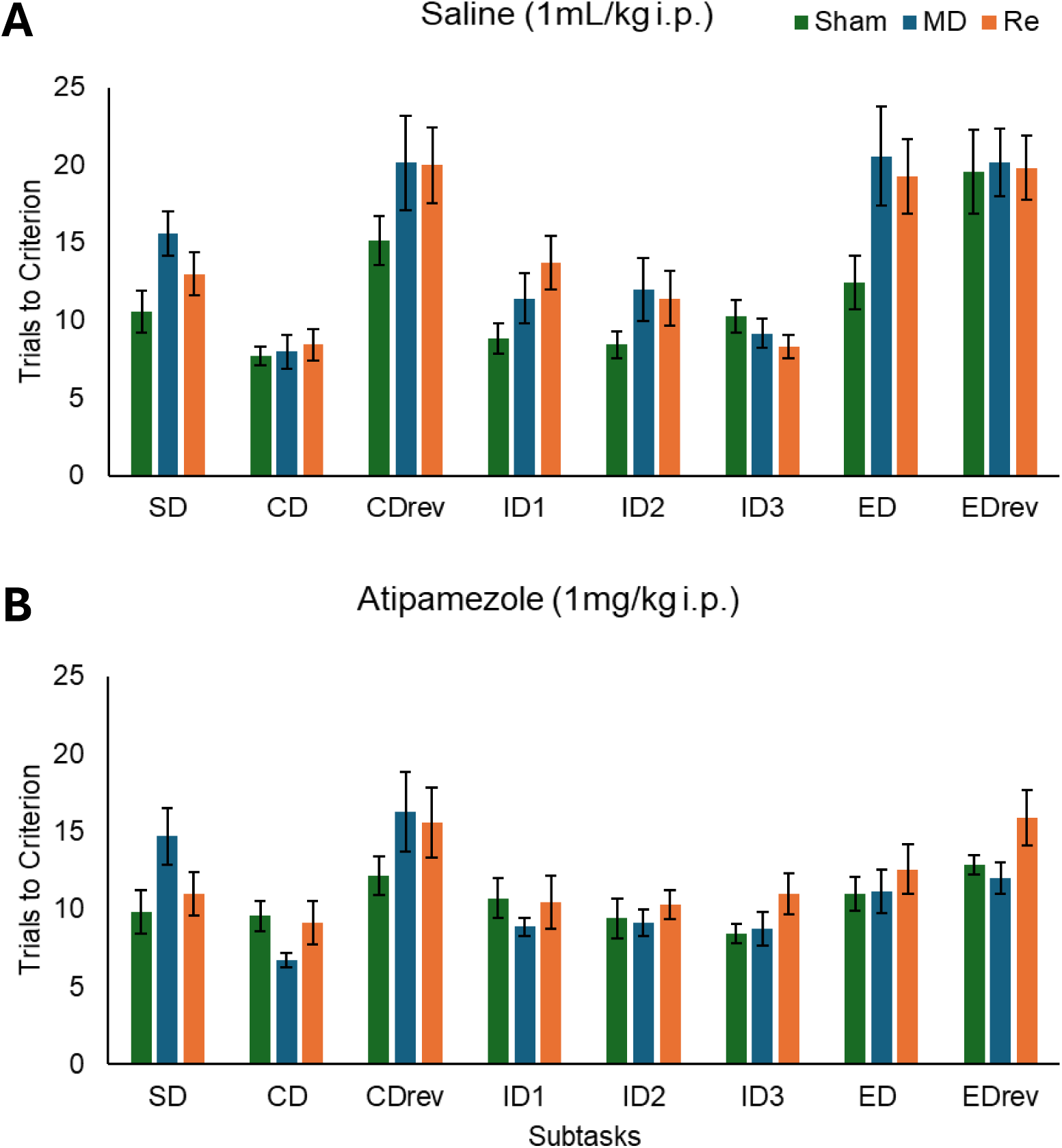
Mean (+/-SEM) for Trials to Criterion to complete each of the 8 subtasks for the three experimental groups: Sham control neurosurgery (n=7), or excitotoxic NMDA infusions into the mediodorsal (MD) thalamic nuclei (n=7), or into the thalamic nucleus reuniens (RE; n=7), during two separate test sessions of the intradimensional/ extradimensional attentional set-shifting task after **A.** saline (1mL/kg; IDED1) intraperitoneal (i.p.) administration or after **B.** atipamezole (1mL/kg) i.p. administration. Following saline, MD (*p*=.006) or RE (*p*=.003) damage produced more trials to criterion than Sham controls. RE damage produced more trials to criterion in ID1 shift than Sham controls (*p*=.03). MD rats required more trials to criterion in the ED shift compared to Sham controls (*p*<.03). Atipamezole reduced the trials to criterion across all rats compared to saline, improving cognitive flexibility performance even after permanent MD or RE thalamic damage. Abbreviations: CD = compound discrimination; CDrev = reversal of compound discrimination; ED =extradimensional shift; EDrev = reversal of extradimensional shift; ID1 = the first intradimensional shift; ID2 = the second intradimensional shift; ID3 = the third intradimensional shift; SD = simple discrimination.

### Group effects on the SD, CD, and CDrev Cluster - IDED1

The comparison of Group x Cluster revealed no main effect of Group, *F*(2,18)=3.28, *p*=.06, although MD and RE groups required more trials to criterion in each subtask. A main effect of Cluster, *F*(1,18)=14.92, *p*=.001, showed all rats required more trials to complete the CD reversal. There was no Group x Cluster interaction, *F*(2,18)=1.36, *p*=.28.

### Group effects on ID1, ID2, ID3 Cluster - IDED1

Across the three ID subtasks, a Group x Cluster interaction, *F*(2,18)=3.56, *p*=.05 (see Fig. 3A), revealed RE compared to Sham rats required more trials to criterion on ID1 (*p*=.03). Across the remaining two ID subtasks, RE and Sham rats’ trials to criterion were similar (ID2 *p*=.20; ID3 *p=.14*). RE rats demonstrated learning with by fewer trials to criterion required across the completion of ID1, ID2, ID3 (*p*=.008). As expected, MD and Sham rats did not differ across the ID Cluster (*p*>.05). There was no main effect of Group, *F*(2,18)=2.13, *p*=.14 or ID Cluster, *F*(1,2)=3.98, *p*=.06.

### Group effects on the ID3-ED Cluster - IDED1

A significant Group x ID3-ED Cluster interaction, *F*(2,18)=3.69, *p*=.04 revealed that, as expected, MD compared to Sham rats require a greater number of trials to criterion (*p*=.03; see Fig. 3A) indicating a deficit in shifting attention to the previously ignored sensory dimension. RE rats did not differ to either Sham (*p*=.06) or MD rats (*p*=.72). All rats performed similarly on ID3 (*p*’s>.10). There was a main effect of ID3-ED Cluster, *F*(1,18)=27.07, *p*<.001, with all rats requiring more trials to complete the ED subtask compared to ID3. There was no Group main effect, *F*(2,18)=1.86, *p*=.18.

### Group effect on ED subtask - IDED1

While MD or RE damaged rats required more trials to criterion than Shams on the ED subtask (Fig. 3A), univariate ANOVA (Welch’s) showed only a non-significant trend for the Group main effect, *F*(2,11.3)=3.85, *p*=.053.

### Group effects on the reversal subtasks (CDrev and EDrev) - IDED1

All rats that completed both reversals (Sham=7; MD=6; RE=6) required a similar number of trials to criterion for the CDrev and EDrev subtasks with no main effect of Group, *F*(2,16)<1.0; no main effect of Subtask, *F*(1,16)=1.18, *p*=.29, and no interaction, *F*(2,16)<1.0. *Atipamezole Group effects on overall IDED2*

Analysis of all 8 subtasks (SD-EDrev) on IDED2 revealed a Group x Subtask interaction, *F*(2,18)=4.62, *p*=.04; see Fig. 3B), a main effect of Subtask, *F*(1,18)=6.07, *p*=.02), but no main effect of Group, *F*(2,18)=2.72, *p*=.09. The significant effects are described below.

### Atipamezole Group effects on the SD, CD, and CDrev Cluster - IDED2

A Group x Cluster interaction, *F*(2,18)=4.77, *p*<.02 revealed that MD rats required significantly more trials to criterion to learn the SD (Fig. 3B; MD vs Sham, *p*=.04), but once the SD was acquired these rats were no different to the RE or Sham rats (CD and CDrev *p*’s>.062). There was a main effect of Cluster, *F*(1,18)=23.71, *p*<.001, with all rats requiring more trials to criterion to complete the CDrev. There was no main effect of Group, *F*(2,18)=1.56, *p*=.23.

### Atipamezole Group effects on the ID1, ID2, ID3 Cluster - IDED2

Analysis of the ID Cluster revealed no main effect of Group, *F*(2,18)=2.19, *p*=.14, or Cluster, *F*(1,18)<1.0, and no Group x Cluster interaction, *F*(2,18)=7.73, *p*=.59, suggesting that atipamezole mitigated the ID1 subtask learning deficit observed during IDED1 in the RE rats (Fig. 3B).

### Atipamezole Group effects on ID3, ED Cluster - IDED2

This analysis showed a main effect of Group, *F*(2,18)=4.29, *p*=.03 (see Fig. 3B) with RE rats requiring more trials to criterion to complete this Cluster than Sham rats (*p*=.05) but no other group differed (*p*’s>.05). There was no main effect of Cluster, *F*(1,18)=3.09, *p*=.09, and no Group x Cluster interaction, *F*<1.0.

### Atipamezole Group effects on ED subtask - IDED2

The analysis showed no main effect of Group, *F*<1.0, with all rats requiring a similar number of trials to complete this ED shift, suggesting that atipamezole had mitigated the ED shift performance observed in IDED1 (Fig. 3B).

### Atipamezole Group effects on the CDrev and EDrev Cluster - IDED2

All rats had similar numbers of trials to criterion for the CDrev and EDrev subtasks (Fig. 3B), with no difference for Group, *F*(2,18)=1.44, *p*=.26, or for Subtask, *F*<1.0, and no Group x Subtask interaction, *F*(2,18)=1.69, *p*=.21.

*Overall Group Effects of Drug Administration: Saline (IDED1) vs Atipamezole (IDED2)* The repeated measures ANOVA (Group x Drug Session x Subtask; Fig. 3) revealed that the Group main effect was still apparent with both MD or RE damaged groups of rats requiring more trials to criterion compared to Sham controls, *F*(2,16)=5.77, *p*=.01; MD vs Sham, *p*=.02; Sham vs RE, *p*=.006; MD vs RE, *p*=.57. Notably, administration of atipamezole prior to IDED2 significantly reduced the number of trials to criterion (Drug main effect, *F*(1,16)=28.35, *p*<.001; saline (M)=13.30, SD=0.89; atipamezole (M)=11.09, SD=0.77). However, there was no Group x Drug interaction, *F*(2,16)=3.01, *p*=.07. The main effect of Subtask remained when combined across the two test sessions, *F*(1,16)=17.42, *p*<.001.

### Saline vs atipamezole Group effects on SD, CD, CDrev cluster

A Group x Cluster interaction [order5], *F*(2,16)=6.04, *p*=.01, indicated that MD rats specifically required more trials to criterion to acquire the SD subtask irrespective of Drug Session compared to Sham control (*p*=.004) or RE (*p*=.03) rats. MD rats also required more trials to criterion for the CDrev subtask than Sham rats (*p*=0.04). There was also a main effect of Group *F*(2,18)=4.43, *p*=.02.

### Saline vs atipamezole Group effects on ID1, ID2, ID3 cluster

Despite the RE deficit following saline administration (IDED1), there were no Group differences or interactions between the saline and atipamezole test sessions for the ID1, ID2 and ID3 subtasks (*p*’s>.05), indicating that atipamezole improved cognitive performance.

### Saline vs atipamezole Group effects on ED subtask

A significant effect of Drug Session *F*(1,18)=12.55, *p*=.002, on the ED subtask shows atipamezole reduced the number of trials to criterion on the ED subtask compared to saline administration. There was no main effect of Group *F*(2,18)=2.87, *p*=.08, and no Group x Drug session interaction *F*(2,18)=2.01, *p*=.16.

### Saline vs atipamezole Group effects on the Shift Cost (ID3-ED)

A significant main effect of Drug Session, *F*(1,18)=10.05, *p*=.005, indicates a decrease in Shift Cost following atipamezole injections compared with saline (see Fig. 4). This drug effect was consistent across all three groups (Group x Drug Session, *F*(2,18)=2.89, *p*=.08), with no main effect of Group, *F*(2,18)=1.82, *p*=.19. Pairwise comparisons showed that atipamezole did not alter responding on the ID3 subtask (*p*=.85), but it significantly reduced trials to criterion on the ED subtask (*p*=.002).

**Figure 4.**
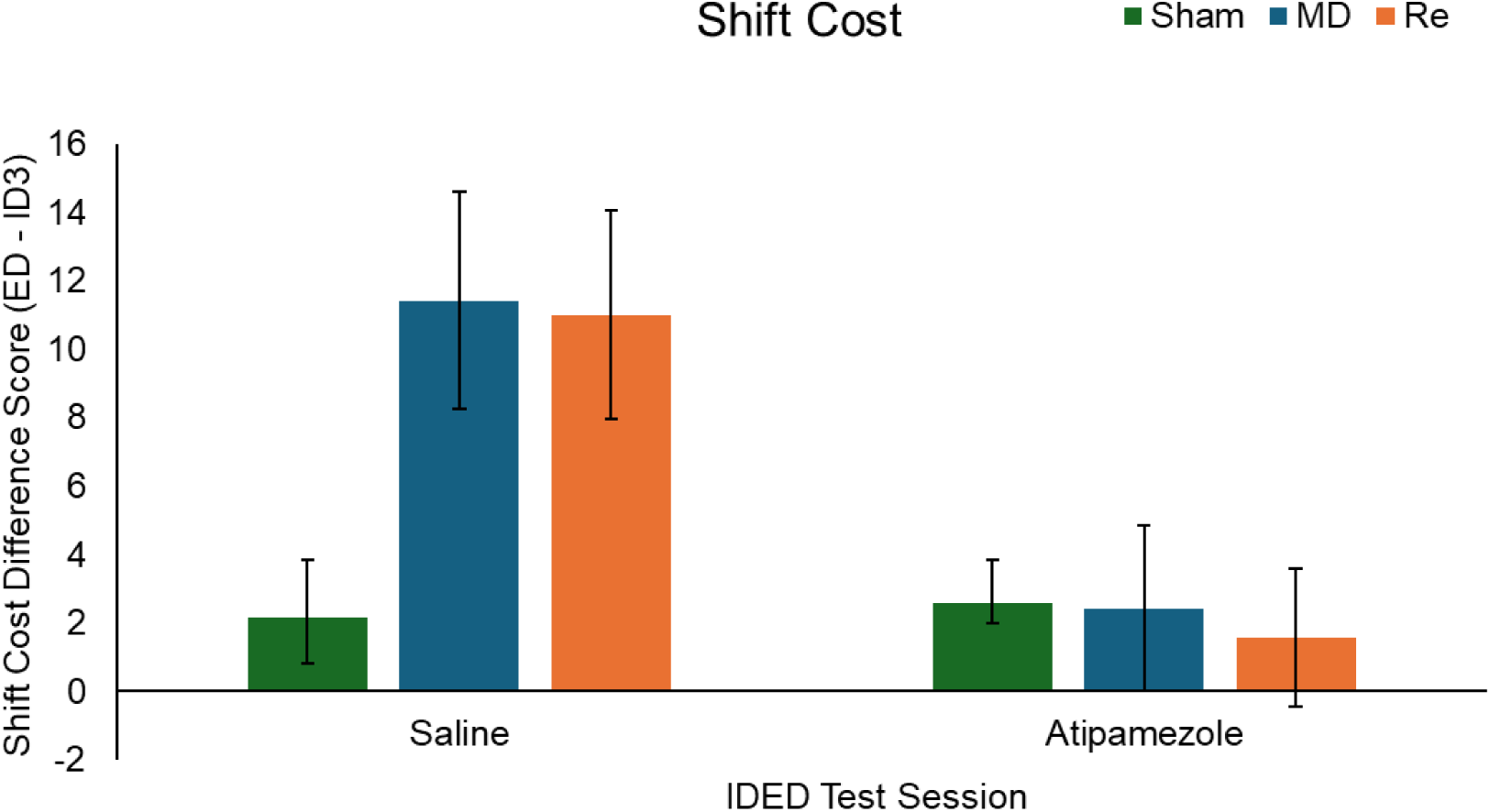
Mean (+/-SEM) Difference Score (ED - ID3) for trials to criterion for the three experimental groups: Sham control neurosurgery (n=7), or excitotoxic NMDA infusions into the mediodorsal (MD) thalamic nuclei (n=7) or into the thalamic nucleus reuniens (RE; n=7) during the two test sessions of the intradimensional/ extradimensional attentional set-shifting task after saline (IDED1, 1mL/kg) or after atipamezole (IDED2, 1mg/kg) intraperitoneal (i.p.) administration.

### Rapid adaptability across all subtasks (saline vs atipamezole)

To determine whether MD or RE damage altered inhibitory responding across the two drug conditions, the percentage of correct 2^nd^ choice responses for all trials was calculated (2^nd^ choice correct trials/total correct trials). Neither MD nor RE damage impaired adaptability across either session (Group main effect, *F*(1,18)=2.34, *p*=0.125; Group x Drug Session interaction, *F*<1.0). Adaptability scores did differ significantly across the two sessions, with a significantly lower score apparent following atipamezole administration (Drug Session main effect, *F*(1,18)=12.61, *p*=.002, possibly due to the significantly reduced number of 2^nd^ entries following atipamezole compared to saline, Drug Session main effect, *F*(1,18)=17.22, *p*<.001; Saline M=24.04, Atipamezole M=12.67).

### Group Average Latencies across all subtasks for saline vs atipamezole

Comparison of the average latencies to complete all trials during the two IDED tests revealed a main effect of Drug Session, *F*(1,18)=23.51, *p*<0.001, with the latency to respond reduced after atipamezole compared to saline, but there was no main effect of Group, *F*<1.0, and no interaction, *F*<1.0.

Examination of general activity in an open field following atipamezole is presented in Supplementary Materials.

## Discussion

The present study showed using the attentional set-shifting IDED task that, as predicted, permanent excitotoxic damage to glutamatergic neurons in the MD impairs the ED shift [9], while permanent excitotoxic damage to glutamatergic neurons in the RE impairs the first ID shift [13], but interestingly RE damage did not impair the additional two ID shift subtasks (ID2 and ID3). This result demonstrates for the first time that rats with permanent RE damage show dissociable performance from rats with permanent excitotoxic anterior thalamic nuclei damage, whereby the RE supports the initial learning of novel ID pairs (ID1) but not the ongoing implementing of the ID set-shifting strategy across the two further ID shift subtasks. In contrast, the anterior thalamic nuclei support both the initial learning of novel ID pairs and the implementing of this set-shifting strategy [40]. Further, the MD does not impact on either component of the ID subtask shift performance.

Additionally, we provide the first evidence that the selective ⍺2-adrenoceptor antagonist, atipamezole administered 30-min before IDED re-testing, reduces the deficits in cognitive flexibility caused by permanent MD or RE damage. That is, across the attentional set-shifting task, atipamezole administration, which is known to increase the release of norepinephrine centrally in the brain [34], reduced the number of trials to criterion, irrespective of thalamic damage, indicating a global facilitation on task performance, rather than a selective rescue of damage related deficits. Consistent with this, some permanent effects persisted across the saline vs atipamezole drug sessions, with MD or RE damaged rats still requiring more trials to criterion overall than Sham controls. Nonetheless, the benefits of using this selective ⍺2-adrenoceptor antagonist were evident, especially during the extradimensional (ED) shift and its reversal, aligning with previous evidence that atipamezole preferentially enhances ED shift cognitive flexibility on the attentional set-shifting task [26,29].

Across the IDED task, atipamezole administration produced robust effects on behavioural performance. Latencies to complete the trials were significantly reduced compared with saline, reflecting faster and more decisive responding. This is consistent with evidence that increased norepinephrine enhances attentional focus and efficiency of task-specific decision making [45], in line with its established role in regulating arousal and effort allocation [26,29,36]. Atipamezole also reduced second-choice entries, suggesting it supports more decisive choice selection. Together, these findings indicate that atipamezole promotes rapid, efficient, and focussed decision-making across the IDED subtasks.

The dissociable effects of MD and RE lesions in cognitive performance across the IDED are consistent with prior evidence that these midline thalamic nuclei support distinct roles in attention, learning and decision-making. The selective impairment following MD damage during the ED shift, while sparing performance on the ID shift, aligns with evidence that the MD is critical for updating attentional set-shifting to previously irrelevant stimuli by rapidly supporting the reconfiguration of choice responses when task rules change unpredictably. Additionally, MD damage also affected the initial learning in the SD and the CDRev after saline administration, while after atipamezole administration acquisition of the new SD stimulus pairing continued to be disrupted. This supports previous evidence gathered across mice, rats, monkeys and humans using differing tasks that each assess new learning and cognitive flexibility [2,7,9,10,18]. The evidence indicates that MD thalamocortical interactions with the mPFC are crucial for cognitive flexibility because when damaged rats produce selective deficits during the SD and ED shift [21,22,25].

One of the underlying mechanisms leading to these deficits may be attributed to the disrupted convergence of signals within the mPFC, including from the hippocampus, the MD, and the ventral tegmental area as previously evidenced [46]. Our immunohistochemistry results demonstrated significantly increased vGluT2 labelling in area 24a ACC of the rodent mPFC after permanent NMDA excitotoxic MD damage compared to the Sham control rats suggesting hyper-functionality in this key mPFC brain region. Increased expression of vGluT2 in other brain areas is shown to produce over compensatory responses and re-modelling of synapses [47]. While additional evidence is still required to further decipher the changes in mPFC, our evidence of impaired cognitive flexibility, especially during the ED shift after loss of MD thalamocortical inputs to the mPFC also mimics the effects of noradrenergic deafferentation within the mPFC or damage to the dorsal noradrenergic bundle that innervates the mPFC [37,38]. Thus our findings implicate the importance of the noradrenergic system in cognition [29] and together they provide evidence that links MD thalamocortical interactions and the noradrenergic system for cognition flexibility [48].

In contrast, RE damaged rats showed deficits during the first ID shift but then comparable performance to Sham controls across the other two ID subtasks. This new evidence is consistent with proposals that the RE supports strategy learning, potentially facilitating the convergence of hippocampal-mPFC signals required for rule formation [11]. Intriguingly, vGluT2 labelling in area 24a ACC was significantly reduced after permanent NMDA excitotoxic RE damage compared to the MD and the Sham rats, suggesting ongoing hypo-functionality in the mPFC as a consequence of the loss of the RE thalamic input. While both the MD and RE project to the mPFC including Layers II-III of ACC, area 24a [49–51], we observed dissociable VGluT2 expression after the permanent thalamic neuron loss. Together, this evidence indicates dissociable findings in behavioural performance and immunohistology whereby the mPFC shows both long-term hyper- and hypo-structural changes and that MD-PFC inputs primarily facilitate rapid updating of task-relevant choice responding, while RE-PFC inputs contribute to the formation of strategies used in this IDED task.

Notably, baseline response latencies (IDED1/saline) were longer than typically reported in the literature. Previous studies employing the attentional set-shifting task in rats have used other strains (Long-Evans and Sprague-Dawley), that generally show faster response times and greater exploration in operant or digging-based tasks [25,52]. In contrast, research has shown that PVGc rats, are known to exhibit lower overall locomotion and more cautious behaviour [53,54], potentially contributing to the slower response times observed here. Despite this, the overall pattern of results aligns with previous reports. There were no lesion-related differences in average latency across IDED1 suggesting similar task engagement across the groups. In contrast, atipamezole administration leads to increased release of norepinephrine and consistently reduced these response latencies, demonstrating that the task remained sensitive to neuromodulatory effects. This highlights that strain differences can influence performance measures without affecting the underlying cognitive processes or pharmacological responsiveness and emphasises the importance of strain characteristics when comparing behaviour across studies.

A limitation of this study is the reliance on systemic pharmacology, which does not isolate noradrenergic mechanisms at specific receptor subtypes or neuroanatomical loci. Future work using regionally targeted manipulations will be required to dissect how noradrenergic signalling interacts with thalamocortical pathways during set shifting. Additionally, electrophysiological or imaging markers of thalamocortical communication during task performance may help clarify the dynamic MD or RE contributions to rule learning and updating.

To conclude, the mPFC was altered after permanent MD or RE damage as evidenced by dissociable VGluT2 expression in the area 24a ACC. Permanent damage to the MD or RE produced dissociable cognitive deficits in the attentional set-shifting task. Specifically, RE damage disrupted learning the first ID shift but the ED shift remained relatively intact. In contrast, MD damage disrupted the ED shift but left ID shifts intact. Atipamezole i.p. injections improved cognitive flexibility for all rats, reducing trials to criterion in all subtasks highlighting the potential therapeutic efficacy of this selective ⍺2-adrenoceptor antagonist to facilitate cognitive flexibility.

## Data Availability Statement

Data will be available on request.

## Author contributions

JJH, JCDA and ASM: funding, concept and design, writing and editing drafts, statistical analysis and interpretation; JJH, LB and SH-B: conducting experiment.

## References

1. Hwang, K., et al., Neuropsychological evidence of multi-domain network hubs in the human thalamus. Elife, 2021. 10: p. e69480.

2. Wolff, M. and M.M. Halassa, The mediodorsal thalamus in executive control. Neuron, 2024. 112(6): p. 893–908.

3. Perry, B.A., E. Lomi, and A.S. Mitchell, Thalamocortical interactions in cognition and disease: The mediodorsal and anterior thalamic nuclei. Neuroscience & Biobehavioral Reviews, 2021. 130: p. 162–177.

4. Pergola, G., et al., The regulatory role of the human mediodorsal thalamus. Trends in cognitive sciences, 2018. 22(11): p. 1011–1025.

5. Lange, F., C. Seer, and B. Kopp, Cognitive flexibility in neurological disorders: Cognitive components and event-related potentials. Neuroscience & Biobehavioral Reviews, 2017. 83: p. 496–507.

6. Browning, P.G., S. Chakraborty, and A.S. Mitchell, Evidence for mediodorsal thalamus and prefrontal cortex interactions during cognition in macaques. Cerebral cortex, 2015. 25(11): p. 4519–4534.

7. Chakraborty, S., et al., Critical role for the mediodorsal thalamus in permitting rapid reward-guided updating in stochastic reward environments. Elife, 2016. 5: p. e13588.

8. Mitchell, A.S., The mediodorsal thalamus as a higher order thalamic relay nucleus important for learning and decision-making. Neuroscience & Biobehavioral Reviews, 2015. 54: p. 76–88.

9. Ouhaz, Z., et al., Mediodorsal thalamus is critical for updating during extradimensional shifts but not reversals in the attentional set-shifting task. Eneuro, 2022. 9(2).

10. Suthaharan, P., et al., Lesions to the mediodorsal thalamus, but not orbitofrontal cortex, enhance volatility beliefs linked to paranoia. Cell reports, 2024. 43(6).

11. Zhang, X., et al., Mediodorsal thalamus regulates task uncertainty to enable cognitive flexibility. Nature Communications, 2025. 16(1): p. 2640.

12. Rikhye, R.V., A. Gilra, and M.M. Halassa, Thalamic regulation of switching between cortical representations enables cognitive flexibility. Nature neuroscience, 2018. 21(12): p. 1753–1763.

13. Linley, S.B., M.M. Gallo, and R.P. Vertes, Lesions of the ventral midline thalamus produce deficits in reversal learning and attention on an odor texture set shifting task. Brain research, 2016. 1649: p. 110–122.

14. Rojas, A.K., S.B. Linley, and R.P. Vertes, Chemogenetic inactivation of the nucleus reuniens and its projections to the orbital cortex produce deficits on discrete measures of behavioral flexibility in the attentional set-shifting task. Behavioural Brain Research, 2024. 470: p. 115066.

15. Panzer, E., et al., In relentless pursuit of the white whale: A role for the ventral midline thalamus in behavioral flexibility and adaption? Neuroscience & Biobehavioral Reviews, 2024. 163: p. 105762.

16. Hoover, W.B. and R.P. Vertes, Collateral projections from nucleus reuniens of thalamus to hippocampus and medial prefrontal cortex in the rat: a single and double retrograde fluorescent labeling study. Brain Structure and Function, 2012. 217(2): p. 191–209.

17. Varela, C., et al., Anatomical substrates for direct interactions between hippocampus, medial prefrontal cortex, and the thalamic nucleus reuniens. Brain Structure and Function, 2014. 219(3): p. 911–929.

18. Monchi, O., et al., Wisconsin Card Sorting revisited: distinct neural circuits participating in different stages of the task identified by event-related functional magnetic resonance imaging. Journal of Neuroscience, 2001. 21(19): p. 7733–7741.

19. Buchsbaum, B.R., et al., Meta-analysis of neuroimaging studies of the Wisconsin Card-Sorting task and component processes. Human brain mapping, 2005. 25(1): p. 35–45.

20. Nyhus, E. and F. Barceló, The Wisconsin Card Sorting Test and the cognitive assessment of prefrontal executive functions: a critical update. Brain and cognition, 2009. 71(3): p. 437–451.

21. Dias, R., T. Robbins, and A. Roberts, Primate analogue of the Wisconsin Card Sorting Test: effects of excitotoxic lesions of the prefrontal cortex in the marmoset. Behavioral neuroscience, 1996. 110(5): p. 872.

22. Dias, R., T.W. Robbins, and A.C. Roberts, Dissociation in prefrontal cortex of affective and attentional shifts. Nature, 1996. 380(6569): p. 69–72.

23. Nakahara, K., et al., Functional MRI of macaque monkeys performing a cognitive set-shifting task. Science, 2002. 295(5559): p. 1532–1536.

24. Brown, V.J. and D.S. Tait, Attentional set-shifting across species. Current Topics in Behavioral Neuroscience, 2016. 28(11): p. 363–395.

25. Birrell, J.M. and V.J. Brown, Medial frontal cortex mediates perceptual attentional set shifting in the rat. Journal of Neuroscience, 2000. 20(11): p. 4320–4324.

26. Lapiz, M. and D. Morilak, Noradrenergic modulation of cognitive function in rat medial prefrontal cortex as measured by attentional set shifting capability. Neuroscience, 2006. 137(3): p. 1039–1049.

27. Owen, A.M., et al., Extra-dimensional versus intra-dimensional set shifting performance following frontal lobe excisions, temporal lobe excisions or amygdalo-hippocampectomy in man. Neuropsychologia, 1991. 29(10): p. 993–1006.

28. Bubb, E.J., et al., Chemogenetics reveal an anterior cingulate–thalamic pathway for attending to task-relevant information. Cerebral Cortex, 2021. 31(4): p. 2169–2186.

29. Poe, G.R., et al., Locus coeruleus: a new look at the blue spot. Nature Reviews Neuroscience, 2020. 21(11): p. 644–659.

30. Holland, N., T.W. Robbins, and J.B. Rowe, The role of noradrenaline in cognition and cognitive disorders. Brain, 2021. 144(8): p. 2243–2256.

31. Marien, M.R., F.C. Colpaert, and A.C. Rosenquist, Noradrenergic mechanisms in neurodegenerative diseases: a theory. Brain Research Reviews, 2004. 45(1): p. 38–78.

32. Scheinin, M., Lomasney, J. W., Hayden-Hixson, D. M., Schambra, U. B., Caron, M. G., Lefkowitz, R. J., & Fremeau Jr, R. T. (1994). Distribution of α2-adrenergic receptor subtype gene expression in rat brain. Molecular Brain Research, 21(1-2), 133–149.

33. Tavares, A., Handy, D. E., Bogdanova, N. N., Rosene, D. L., & Gavras, H. (1996). Localization of α2A-and α2B-adrenergic receptor subtypes in brain. Hypertension, 27(3), 449–455.

34. Pertovaara, A., Haapalinna, A., Sirviö, J., & Virtanen, R. (2005). Pharmacological properties, central nervous system effects, and potential therapeutic applications of atipamezole, a selective α2-adrenoceptor antagonist. CNS drug reviews, 11(3), 273–288.

35. Scheinin, H., MacDonald, E., & Scheinin, M. (1988). Behavioural and neurochemical effects of antipamezole, a novel α2-adrenoceptor antagonist. European journal of pharmacology, 151(1), 35–42.

36. Haapalinna, A., Viitamaa, T., MacDonald, E. et al. (1997). Evaluation of the effects of a specific α2-adrenoceptor antagonist, atipamezole, on α1- and α2-adrenoceptor subtype binding, brain neurochemistry and behaviour in comparison with yohimbine. Naunyn-Schmiedeberg’s Arch Pharmacol 356, 570–582.

37. Tait, D.S., et al., Lesions of the dorsal noradrenergic bundle impair attentional set-shifting in the rat. European Journal of Neuroscience, 2007. 25(12): p. 3719–3724.

38. McGaughy, J., R.S. Ross, and H. Eichenbaum, Noradrenergic, but not cholinergic, deafferentation of prefrontal cortex impairs attentional set-shifting. Neuroscience, 2008. 153(1): p. 63–71.

39. Janitzky, K., et al., Optogenetic silencing of locus coeruleus activity in mice impairs cognitive flexibility in an attentional set-shifting task. Frontiers in Behavioral Neuroscience, 2015. 9: p. 286.

40. Wright, N.F., et al., A critical role for the anterior thalamus in directing attention to task-relevant stimuli. Journal of Neuroscience, 2015. 35(14): p. 5480–5488.

41. Hamilton, J.J. and J.C. Dalrymple-Alford, Anterior thalamic nuclei: A critical substrate for non-spatial paired-associate memory in rats. European Journal of Neuroscience, 2022. 56(7): p. 5014–5032.

42. Fujiyama, F., Furuta, T. and Kaneko, T. (2001), Immunocytochemical localization of candidates for vesicular glutamate transporters in the rat cerebral cortex. J. Comp. Neurol., 435: 379–387.

43. Herzog, E., Bellenchi, G. C., Gras, C., Bernard, V., Ravassard, P., Bedet, C., … & El Mestikawy, S. (2001). The existence of a second vesicular glutamate transporter specifies subpopulations of glutamatergic neurons. The Journal of Neuroscience, 21(22), RC181.

44. Kaneko T, Fujiyama F. Complementary distribution of vesicular glutamate transporters in the central nervous system. Neurosci Res. 2002 Apr;42(4):243–50.

45. Aston-Jones, G. and J.D. Cohen, An integrative theory of locus coeruleus-norepinephrine function: adaptive gain and optimal performance. Annual Review of Neuroscience, 2005. 28(1): p. 403–450.

46. Floresco, S. B., & Grace, A. A. (2003). Gating of hippocampal-evoked activity in prefrontal cortical neurons by inputs from the mediodorsal thalamus and ventral tegmental area. Journal of Neuroscience, 23(9), 3930–3943.

47. Zeng C, Nannapaneni N, Zhou J, Hughes LF, Shore S. Cochlear damage changes the distribution of vesicular glutamate transporters associated with auditory and nonauditory inputs to the cochlear nucleus. J Neurosci. 2009 Apr 1;29(13):4210–7

48. Shine, J. M., Neuromodulatory control of complex adaptive dynamics in the brain. Interface focus, 2023. 13(3).

49. Krettek, J. E., & Price, J. L. (1977). The cortical projections of the mediodorsal nucleus and adjacent thalamic nuclei in the rat. Journal of comparative neurology, 171(2), 157–191.

50. Wang, C. C., & Shyu, B. C. (2004). Differential projections from the mediodorsal and centrolateral thalamic nuclei to the frontal cortex in rats. Brain research, 995(2), 226–235.

51. Vertes, R. P., Hoover, W. B., Do Valle, A. C., Sherman, A., & Rodriguez, J. J. (2006). Efferent projections of reuniens and rhomboid nuclei of the thalamus in the rat. The Journal of comparative neurology, 499(5), 768–796.

52. Chudasama, Y. and T.W. Robbins, Dissociable contributions of the orbitofrontal and infralimbic cortex to pavlovian autoshaping and discrimination reversal learning: further evidence for the functional heterogeneity of the rodent frontal cortex. Journal of Neuroscience, 2003. 23(25): p. 8771–8780.

53. Broadhurst, P., Determinants of emotionality in the rat: II. Strain differences. Journal of Comparative and Physiological Psychology, 1958. 51(1): p. 55.

54. Schmitt, U. and C. Hiemke, Strain differences in open-field and elevated plus-maze behavior of rats without and with pretest handling. Pharmacology Biochemistry and Behavior, 1998. 59(4): p. 807–811.

