## Supplemental Materials and Methods for "Noradrenergic administration improves cognitive flexibility after permanent damage to glutamatergic neurons in rat mediodorsal thalamus or thalamic nucleus reuniens"

### Supplementary Material

### Materials and Methods

*Animals*

Twenty-four male Piebald Virol Glaxo cArc (PVGc) rats weighing between 315-415g at time of surgery were housed four per cage in a temperature (21±2°C) and humidity (50±5%) controlled holding room. Lights were on a reverse 12-hour light-dark cycle (lights on 8pm), with all behavioural testing completed in the dark (i.e., more active) cycle for the rats. The rats were on a controlled feeding schedule (between 12-18g/rat per day) to maintain 85% of their free-feeding body weight with *ad libitum* access to water in their home cage. Weekly cage cleaning occurred after the rat's behavioural training and testing for that given day. The University of Canterbury Animal Ethics Committee approved all housing and experimental procedures (2022/14R).

*Neurosurgery to create permanent MD or RE thalamic damage*

Rats were pseudorandom assigned to one of three groups (MD, N= 8; RE, N = 8; Sham, N=8). Rats were anaesthetised using isoflurane (4% induction; 2-2.5% maintenance), the shaved head was cleaned with chlorohexidine (Hibitane) using aseptic technique and secured in a stereotaxic frame (Kopf, Tujunga, CA). The incisor bar was set to -5.5mm below interaural line to minimise syringe damage to the fornix. Methopt Forte eye drops (Aspen Pharm, NZ) were administered and a wet gauze pad applied above, but not touching the eyes to stop them drying out and minimise the impact of surgery lights. A midline incision was made, the skull exposed and stereotaxic coordinates made relative to bregma and lambda. To create the permanent damage in either the MD or RE, 0.12M N-methyl-d-aspartate (NMDA; Sigma, Castle Hill, NSW), a potent glutamate receptor agonist, was bilaterally infused with one infusion in each hemisphere of the MD or the RE. For MD infusions, to ensure greater AP spread, one infusion was made more anteriorly in one hemisphere at AP -3.75mm from

Bregma, and more posteriorly in the contralateral hemisphere, at AP -3.95mm from Bregma. Both MD infusions were made ±0.7mm from midline, at a depth of -5.70mm below dura. The infusion volume was 0.20µl NMDA at each MD infusion site at a rate of 0.04µl/min. For RE infusions, bilateral infusions were made at a 10-degree angle -2.3mm from Bregma, ±1.1mm from midline, at a depth of -7.0mm below dura. The infusion volume was 0.12µl NMDA at each RE infusion site at a rate of 0.04µl/min. For each MD or RE infusion, the syringe needle remained in situ at the end of each infusion for 4mins diffusion time. Sham infusion control surgeries used the same surgical procedures and stereotaxic coordinates, but the syringe needle was lowered to 1mm above each of the MD or RE infusion sites and no infusion was administered.

*IDED attentional set-shifting task*

*Apparatus*

The apparatus (**Error! Reference source not found.**A; 40cm x 60cm x 18cm) was placed on a 70cm high table. It contained three compartments made from opaque black Perspex: a large start compartment (40cm x 36cm) and two identical choice compartments (20cm x 24cm) to which access was controlled by clear Perspex doors (20cm x 20cm). Black ceramic digging bowls (7cm diameter x 3.5cm high) with wire mesh epoxy glued ~1cm inside the bottom of the bowl (for hiding an unreachable reward, see *Procedures*) were placed inside each of the two choice compartments. The bowls were filled with clean bedding sawdust scented with odours (herbs or spices), digging media, or a combination of odours and digging media (examples, Fig. 1B).

*Preoperative dig training*

Before surgery, rats maintained at 85% of their free-feeding weight were taught to dig in bowls filled with sawdust to retrieve one half of a Honey Cheerio (Uncle Toby's, Australia). The Honey Cheerio was initially placed on the top of the sawdust and gradually lowered until it was completely hidden from view in subsequent training trials. The Honey Cheerio was rebaited (fully concealed) in each bowl (both choice compartments baited) until the rat made 6 retrievals from each choice compartment. It took up to 3 days and 6 consistent digs in each bowl for all rats to complete dig training.

*IDED Testing Procedures*

The IDED attention set-shifting task is self-paced and relies on the natural foraging behaviours of the rats. During the experiment, one of the two ceramic digging bowls is baited with the reward (Honey Cheerio) while the other one is unbaited, and the rat learns which bowl is baited using either the types of digging media or odours as cues to determine the reward. A hidden reward is placed underneath the wire mesh in each digging bowl to reduce odour cues between baited and non-baited bowls.

At the beginning of each trial, the rat is held in the larger compartment, and the clear Perspex panels are removed to allow the rat access to both the two smaller choice compartments (Fig. 1A; correct choice counterbalanced across right / left). If an incorrect choice is made, this is recorded as an error, and for the first four trials only of any subtask, the rat is allowed access to the correct choice compartment to dig in the bowl to receive the reward before being returned to the larger compartment area and the Perspex panels blocking entry until the start of the next trial (~10-20 second inter trial interval).

For this set of experiments, the IDED task was run twice (IDED1 and IDED2). IDED1 was divided into two sessions comprised of a simple discrimination (SD) training session for the first session and on the next day (see *Postoperative training on the simple discrimination*), the full IDED attentional set-shifting test session. The full IDED attentional set-shifting test session comprised of 8 different subtasks: a novel SD; concurrent discrimination (CD); concurrent discrimination reversal; ID1; ID2; ID3; ED; ED reversal. The correct (rewarded) odour or digging media discriminations and the correct stimulus within pairs (see Figure 1B for stimuli used in IDED1 and IDED2) were counterbalanced across the subtasks and matched between different experimental groups. IDED2 used the same stimuli as IDED1 but the pairings and correct (rewarded) stimulus were shuffled so all pairings were novel to counter any effect of retained learning about a particular stimulus cue between the first (IDED1) and second (IDED2) runs of task. Counterbalancing of the rewarded sensory dimension was also conducted during the SD subtask so that rats who received odour as the rewarded sensory dimension in the IDED1 SD subtask now had digging medium as the rewarded sensory dimension in the IDED2 SD subtask. For IDED2, rats only ran in the full IDED attentional set-shifting test session comprised of the 8 different subtasks, that is, there was no repeat of the initial SD training session prior to running the full test session because the rats already knew how to distinguish between different stimuli in the bowls for rewards linked to specific cues.

*Postoperative training on the simple discrimination*

Rats completed two runs of postoperative testing IDED1 and IDED2 after recovery from surgery. The day before rats completed the full IDED1 session, each rat learnt a simple discrimination (SD) beginning with either odour (all spice versus cardamom in clean bedding sawdust) or digging media (pebbles versus vermiculite) counterbalanced across rats and groups. The rewarded stimulus within any pair was counterbalanced across the groups and rats were trained to a criterion of 6 consecutive correct trials or a maximum of 30 trials per subtask. The stimuli used in the training session were not used again until IDED2. Digging was defined as active digging or foraging with the paws or nose in the digging media, but sniffing or resting paws on the digging media was not counted as an active dig. This digging criterion was used throughout testing on the IDED attentional set shifting task.

*Postoperative testing on IDED attention set-shifting task*

On the next day, the rat was given a series of eight subtasks, each subtask consisted of a series of two-choice trials with only one of the two-choice bowls baited, counterbalanced across side. The rat must learn each new two-choice discrimination to a performance criterion of 6 correct consecutive responses before moving onto the next subtask. The first subtask was a simple discrimination (SD), which is run per the training day but using a new pairing of either odour or digging media, and these pairings were counterbalanced across groups. If a rat was assigned the odour, they would run the odour-to-media trials for IDED1 and media-to-odour for IDED2. For example, in the odour-to-media order of discriminations rats were initially required to attend to the odour sensory dimension, then switched to digging media in the final two subtasks, ED and EDrev.

Following the SD subtask was the compound discrimination (CD) subtask where the same stimulus from the SD is rewarded, with the addition of a new irrelevant sensory dimension (i.e., if odour was learnt on the SD, a digging media pair of stimuli were added but not required to be attended to, or if digging media was learnt as the sensory dimension on the SD, an odour pair was added but not required to be attended to). The CD subtask is followed by the CD reversal (CDrev) used the same stimuli for the CD but the other stimulus in the pair linked to the relevant sensory dimension is now rewarded and the previously rewarded stimulus is the non-rewarded stimulus. This reversal was followed by three separate intradimensional (ID1, ID2, ID3) shift subtasks. For the ID task, the sensory dimension used in the SD and CD subtasks must continue to be attended to in order to receive the rewards, but new stimuli pairings were used for each of the three different ID subtasks. As indicated in the Introduction, previous evidence showed that after damage to the RE, rats require more trials to complete the ID subtask (see [11]) suggesting a deficit in shifting the attentional set strategy after RE damage. So, in the current study, we included two additional ID subtasks, as described previously [40] to determine if damage to the RE disrupts transferring the

In-Confidence

attentional set strategy to novel exemplars, as previously identified after permanent anterior thalamic nuclei damage [40].

For the last two subtasks, extradimensional (ED) shift followed by a reversal of this ED (EDrev), the rats had to rapidly switch their responses to the previously irrelevant non-rewarded sensory dimension after a change in the current task demands. That is, until the end of the three ID shift subtasks, the rats were rewarded if they were digging in the correct bowl of new stimuli pairings linked to the initial sensory dimension (odour or digging media) they learnt from the SD subtask, while ignoring the irrelevant, non-rewarded sensory dimension that was introduced in the CD subtask. Now, for the ED shift, the rats had to learn that they needed to attend to this previously irrelevant sensory dimension (based on negative feedback, i.e., the absence of reward) and learn which stimulus of the new pairing of the now relevant sensory dimension were rewarded. For example, if the rats had been rewarded for attending to the stimuli pairing linked to the odour sensory dimension throughout SD, CD, CDrev, ID1, ID2 and ID3, they had to switch to attend to the previously irrelevant sensory dimension, digging media, and learn to respond to the sensory dimension – touch, while digging in the bowls and determine which digging media stimulus provides reward. Similarly, if the rats had been attending to the sensory dimension of digging media, then they had to switch to attend to the sensory dimension of olfaction and respond to the different odour pairings and determine which stimulus provides reward. The last subtask, the EDrev, involves a reversal of reward contingencies for the stimulus pairing linked to the relevant ED sensory dimension.

*Drug administration*

Prior to running in the first IDED test session (IDED1), each rat received an i.p. injection of saline (1ml/kg) and was placed back into a holding cage for 30 minutes. IDED1 was used to compare cognitive flexibility performance between the three experimental groups (Sham lesion controls, MD lesion or RE lesion). This drug administration procedure was repeated prior to running in the second IDED test session (IDED2) but now with an i.p. injection of atipamezole (1mg/kg, Abcam; [26]) 30 minutes prior to testing. IDED2 was used to determine if atipamezole injection could mitigate any performance deficits in cognitive flexibility caused by MD or RE permanent damage compared to Sham controls.

*Histology*

*Perfusion and tissue collection*

Following behavioural testing, rats were deeply anaesthetised with sodium pentobarbital (125 mg/kg) and perfused transcardially with saline and the brain fixed with paraformaldehyde (4% PFA in 0.1M phosphate buffer; PB). Brains were post-fixed in 4% PFA followed by a minimum of 48 hours in a long-term solution (30% glycerol in 0.1M PB). Coronal 40µm sections were collected using a freezing microtome (Bright Instruments, UK) and stored in cryo-protectant (30% glycerol, 30% ethylene glycol in 0.1M PB) at -20˚C until processed for immunohistochemistry. Coronal sections were collected consecutively in 10 cryovials for immunohistochemistry, with thalamic sections (between -1.08mm and -3.96mm from Bregma) from vials 1, 5, and 9 (i.e., every 160 microns) used to determine the loss of NeuN-stained neurons for verification of the damage.

*NeuN thalamic damage verification*

Per Hamilton & Dalrymple-Alford [41], sections throughout the thalamus were washed in 0.1M phosphate buffered saline with Triton-X (0.2%; PBSTx) before incubation in endogenous peroxidase blocking buffer for 30 minutes (1% hydrogen peroxide ($H_2O_2$), 50% methanol ($CH_3OH$) in 2% PBSTx). They were incubated overnight at 4˚C in anti-NeuN primary antibody (1:5000; monoclonal-Mouse Cat# MAB377; Millipore, California USA) in PBSTx with 1% normal goat serum (NGS; Life Technologies, NZ), followed by washes in PBSTx and incubation in biotinylated goat anti-mouse secondary antibody (1:1000 Cat# BP-9200-50: Vector Laboratories, California USA) for 90min at room temperature in PBSTx and

1% NGS. Sections were placed in ExtrAvidin (peroxidase conjugated; 1:1000; Sigma, NSW Australia), PBSTx and 1% NGS for 2 hours at room temperature. Sections were visualised using diaminobenzidine (DAB 0.05%; Sigma, in 0.01% $H_2O_2$ in Tris buffer; approximately 5 min). After mounting on gelatinised slides, sections were dried and slides were dehydrated through graded alcohol (70-100%), cleared in xylene and mounted with DPX (06522; Sigma Aldrich).

To visualise permanent damage to either the MD or RE, Neu-N positive sections mapped onto atlas plates to determine the extent of each lesion. This used sections from both the left and right hemisphere for every one in four 40μm sections. NeuN-positive cell staining was photographed using the 5x objective on a Leica DM6 B upright microscope and DFC7000T camera (Leica Microsystems, Germany). The area surrounding the thalamic region of interest was manually selected and area recorded using ImageJ, image analysis software (National Institute of Health, NIH, USA) and the percentage of lesion was calculated (area of MD or RE damage / area of MD or RE intact) x 100). Acceptable lesions in the MD or RE were defined as reaching the criterion of at least 60% damage in the respective thalamic region.

*VGluT2 immunofluorescence staining and visualisation*

Thalamocortical projections are glutamatergic with thalamocortical axon terminals defined by the protein, vesicular glutamate transporter 2 (VGluT2 [42-44). To help capture changes in the mPFC after permanent removal of glutamatergic neurons in the MD or RE, we stained sections involving Layer II-III anterior cingulate cortex (ACC) area 24a with VGluT2 protein. Sections from vial 2 were washed in 0.1M phosphate buffered saline with Triton-X (0.2%; PBSTx), non-specific binding was blocked by incubation using 20% NGS 0.2% PBSTx for 60 minutes, then sections were incubated overnight at 4˚C in anti-VGluT2 primary antibody (1:1000; AB216463, Abcam, New Zealand) in PBSTx with 1% NGS. The next day, after further washes in PBSTx, visualisation occurred by incubation in AlexaFluor 488 goat anti-rabbit secondary antibody (1:500, Invitrogen, Oregon, USA) for 90min at room temperature in PBSTx and 1% NGS. After mounting with DAPI (Vectashield antifade mounting medium, Vector Laboratories, Burlingame, CA, USA) to enable nuclear cell visualisation, slides were cover slipped and stored at 4°C in the dark.

The number of VGluT2 terminal puncta present in the sections was used to determine long-term changes to thalamocortical projections following the NMDA excitotoxic damage. VGluT2 and DAPI positive staining was visualised and imaged using the 40x objective with excitation from the L5 (green; LED 470; excitation, 470/40; emission, 525/50; Leica) and UV 'A' (blue, LED 365; excitation, 360/40; emission, long pass 425; Leica) filters respectively. Automated counts of the VGluT2 puncta were obtained through ImageJ. In each image, the two green and blue channels were separated and Layer II and Layer III (individually) of the anterior cingulate cortex (A24a) at approximately 1.28mm from bregma were manually selected. To reduce background noise, images were filter with a gaussian blur (sigma = 1) and background was subtracted (rolling = 60) before converting to mask. Finally, the watershed function was applied, and all puncta above threshold ('Otsu' threshold, circularity 0.2-1.0) were counted.

*Statistical analysis*

The criterion for completing each subtask was 6 correct consecutive trials. Repeated measures ANOVAs evaluated the number of trials to criterion for each rat for each the 8 subtasks (SD, CD, CDrev, ID1, ID2, ID3, ED, EDrev) with Group as the between subject factor (MD, RE or Sham group) for the IDED1 test session when saline i.p. was given prior to testing. The same analysis was conducted for the IDED2 test session, when atipamezole was administered i.p. prior to testing. In the IDED1 test with saline, two rats (one from the MD group and one from the RE group) did not complete the EDrev subtask because they had spent 3 hours completing the first 7 subtasks. For the IDED1 ANOVA the first 7 subtasks were examined.

Due to the complex nature of the interaction between Subtask and Group, subsequent analyses were run to analyse MD or RE thalamic damage effects separately compared to Sham controls for each subtask. The 8 subtasks were divided further into four subtask clusters based on a priori hypotheses of expected deficits following MD or RE damage: Subtask cluster 1= SD, CD, CDrev; Subtask cluster 2= ID1, ID2, ID3; Subtask cluster 3= ID3, ED; and Subtask cluster 4 = ED. Subtask clusters 1, 2 and 3 were analysed using repeated measures ANOVAs (e.g., Subtask 1 [SD, CD, CDrev] x Group) with post-hoc analyses for relevant Subtask cluster x Group effects (2-way ANOVA). Subtask cluster 4 was analysed using a univariate ANOVA with pairwise comparisons for relevant Group effects. Further analyses were conducted for the Shift Cost, using a difference score (ED minus ID3) which compares the trials to criterion of ID3 with the trials to criterion for ED across the Groups. This Shift Cost analysis can provide insight into the behavioural performance of the rats when they need to switch their responses to the previously irrelevant sensory dimension. Additionally, four repeated measures ANOVAs compared trials to criterion for Subtask cluster 1 or Subtask cluster 2 or the ID-ED Shift Cost or the ED subtask across Drug Sessions (IDED1 (saline) x IDED2 (atipamezole)) x Group. Similarly, average latency to complete all trials for Drug Session (IDED1 vs IDED2) were compared using a repeated measures ANOVA (Drug Session x Group). Finally, a repeated measures ANOVA compared differences between Groups of second entry choice responses made for all subtasks combined as a measure of rapid adaptability in choice making.

**Open field investigation of atipamezole related behavioural response**

*Open field exploration pre and post the atipamezole injection*

For five minutes prior to the injection of atipamezole and running the IDED2 session, the rat was placed in a familiar grey Perspex box (60cm x 60cm x 30cm; 30cm above ground level) for 5mins to record baseline (no drug) activity. An overhead camera captured the rat's behaviour for offline analysis. Upon completion of the IDED2 test session, each rat was returned to this open field for another 5mins to record drug activity at about 2-2.5 hours post the atipamezole injection (the delay varied according to the time it took each rat to complete the 8 subtasks in the IDED2 test session).

*General activity (distance travelled) in the open field box*

The repeated measures ANOVA of distance travelled in the open field box prior to the atipamezole administration and immediately on completion of the EDrev subtask showed a significant main effect Pre vs Post injection, $F(1,18) = 18.37$, $p < .001$, with an increase in distance travelled at the end of the EDrev compared with before atipamezole. However, there was no main effect of Group, $F < 1.0$, and no interaction, $F < 1.0$, indicating that the effect of atipamezole on distance travelled by all rats was similar regardless of Group.
